# The effects of transcranial direct current stimulation on corticospinal and cortico-cortical excitability and response variability: conventional versus high-definition montages

**DOI:** 10.1101/2020.03.30.017046

**Authors:** Michael Pellegrini, Maryam Zoghi, Shapour Jaberzadeh

## Abstract

Response variability following transcranial direct current stimulation (tDCS) highlights need for exploring different tDCS electrode montages. This study compared corticospinal excitability (CSE), cortico-cortical excitability and intra-individual variability following conventional and HD anodal (a-tDCS) and cathodal (c-tDCS) tDCS. Fifteen healthy young males attended four sessions at least one-week apart: conventional a-tDCS, conventional c-tDCS, HD-a-tDCS, HD-c-tDCS. TDCS was administered (1mA, 10-minutes) over the primary motor cortex (M1), via 6×4cm active and 7×5cm return electrodes (conventional tDCS) and 4×1 ring-electrodes 3.5cm apart in ring formation around M1 (HD-tDCS). For CSE, twenty-five single-pulse transcranial magnetic stimulation (TMS) peak-to-peak motor evoked potentials (MEP) were recorded at baseline, 0-minutes and 30-minutes post-tDCS. For cortico-cortical excitability, twenty-five paired-pulse MEPs with 3-millisecond (ms) inter-pulse interval (IPI) and twenty-five at 10ms assessed short-interval intracortical inhibition (SICI) and intracortical facilitation (ICF) respectively. MEP standardised z-values standard deviations represented intra-individual variability. No significant differences were reported in CSE between conventional and HD a-tDCS, but significant differences between conventional and HD c-tDCS 0-minutes post-tDCS. Intra-individual variability was significantly reduced in conventional tDCS compared to HD-tDCS for a-tDCS (0-minutes) and c-tDCS (30-minutes). No significant changes were reported in SICI and ICF. These novel findings highlight current technical issues with HD-tDCS, suggesting future tDCS studies should utilise conventional tDCS to minimise intra-individual variability, ensuring tDCS after-effects are true changes in CSE and cortico-cortical excitability.

## Introduction

Transcranial direct current stimulation (tDCS) has risen in popularity in recent years as a tool for inducing cortical plasticity in real-time. As a form of non-invasive brain stimulation (NIBS), it alters the excitability of cortical regions by influencing the underlying neurones membrane potential, evoking either increases or reductions in corticospinal excitability (CSE) and cortico-cortical excitability (i.e. intracortical inhibition and intracortical facilitation) (M. A. Nitsche et al., 2003, 2007). Traditionally delivered via two rectangular surface electrodes, resultant changes in CSE and cortico-cortical excitability are dependent on the electrode polarity (M. A. Nitsche & Paulus, 2000a). Neuronal depolarization and resultant increases in CSE occur when the anode is placed over the cortical area of interest, while hyperpolarization and reductions in CSE occur when the cathode is placed over the cortical area of interest (M. A. Nitsche & Paulus, 2000a). This predictability, along with minimal side-effects and cost-effectiveness, has contributed to its popularity for investigating properties of cortical plasticity in healthy and neurological populations (Chew et al., 2015; Strube et al., 2015).

Over the past decade however, its predictability to induce polarity-dependent changes in CSE has been questioned. Recent studies have reported subgroups of individuals responding to tDCS in a historically polarity-dependent manner and those that do not (Ammann et al., 2017; Chew et al., 2015; López-Alonso et al., 2014; Strube et al., 2015; Wiethoff et al., 2014). Comprehensively reviewed elsewhere (Pellegrini et al., 2018b), termed ‘responders’ and ‘non-responders’, this variability in response to tDCS between individuals, is capable of influencing tDCS overall effects and therefore is a frequently discussed limitation to published tDCS studies when overall significant effects are not achieved (Strube et al., 2015).

Response variability is an umbrella term for inter-individual variability and intra-individual variability. Inter-individual variability, as discussed above, is the variation in response to tDCS between participating individuals. Specific mechanisms to account for inter-individual variability have been discussed previously in recent reviews (Chew et al., 2015; Pellegrini et al., 2018a; Ridding & Ziemann, 2010). Intra-individual variability however refers to the variation in response to tDCS within an individual across different testing sessions. Sources of intra-individual variability include the intra-rater reliability and consistency of the assessor when measuring CSE and cortico-cortical excitability across multiple testing sessions. Additionally, physiological factors such as differences in female responses across different phases of the menstrual cycle (Zoghi et al., 2015), the time-of-day testing sessions are conducted (M. V. Sale et al., 2008; Martin V. Sale et al., 2007), the history of prior synaptic activity (Gentner et al., 2008; Iezzi et al., 2008; Ridding & Ziemann, 2010; Rosenkranz et al., 2007), the level of caffeine intake (Cerqueira et al., 2006; Concerto et al., 2017) or number of hours of sleep prior to testing (Manganotti et al., 2001; Placidi et al., 2013) can influence an individuals’ changes in CSE and cortico-cortical excitability following tDCS.

With response variability commonly discussed as a limitation in tDCS literature, exploration into different forms of tDCS delivery other than conventional tDCS is warranted. A lack of overall significant changes in CSE and cortic-cortical excitability following tDCS despite large cohorts of individuals (Chew et al., 2015; López-Alonso et al., 2014; Puri et al., 2015; Strube et al., 2015, p. 20; Tremblay et al., 2016) calls into question the capacity of conventional tDCS protocols to reliably induce changes in CSE and cortico-cortical excitability and whether reported results are true reflections of changes in CSE and cortico-cortical excitability and not influenced by high response variability.

One different technique is high-definition tDCS (HD-tDCS). The electrode montage in HD-tDCS uses a central active electrode placed directly over the cortical area of interest with four surrounding return electrodes arranged in a ring formation spaced equidistance apart surrounding the central active electrode. Reported to increase the focality of the electric field (EF), a greater magnitude targets the cortical areas of interest rather than dispersed across the scalp as with conventional tDCS (figure 1) (Datta et al., 2008, 2009; Edwards et al., 2013). Magnetic resonance imaging studies have confirmed conventional tDCS and HD-tDCS deliver electric current differently (Datta et al., 2009). With different electrode placement between conventional tDCS and HD-tDCS (Datta et al., 2009) and the large size of the conventional rectangular electrodes (Lang et al., 2005), widespread cortical regions were reported to receive electrical current following conventional tDCS (Datta et al., 2009; Lang et al., 2005) with peak EF not located under the cortical area of interest (figure 1a, 1c) (Datta et al., 2009). HD-tDCS conversely was reported to elicit peak EF magnitude over the cortical area of interest and restricted the majority of the EF to the boundaries of the HD-tDCS return electrodes, minimising spread of EF across the scalp (figure 1b, 1c) (Datta et al., 2009). Given its potential for more targeted delivery of electrical current, the efficacy of HD-tDCS has been investigated in Epilepsy (Karvigh et al., 2017), Fibromyalgia (Castillo-Saavedra et al., 2016; Villamar et al., 2013), Memory (Chua et al., 2017; Hill et al., 2017, 2018; Naka et al., 2018; Nikolin et al., 2015; Price et al., 2016), Pain (Borckardt et al., 2012; Flood et al., 2016) and Tinnitus (Henin et al., 2016; Jacquemin et al., 2018). Despite this, direct comparisons between conventional tDCS and HD-tDCS is limited into their differing effect on CSE, cortico-cortical excitability and response variability.

**Figure 1.**
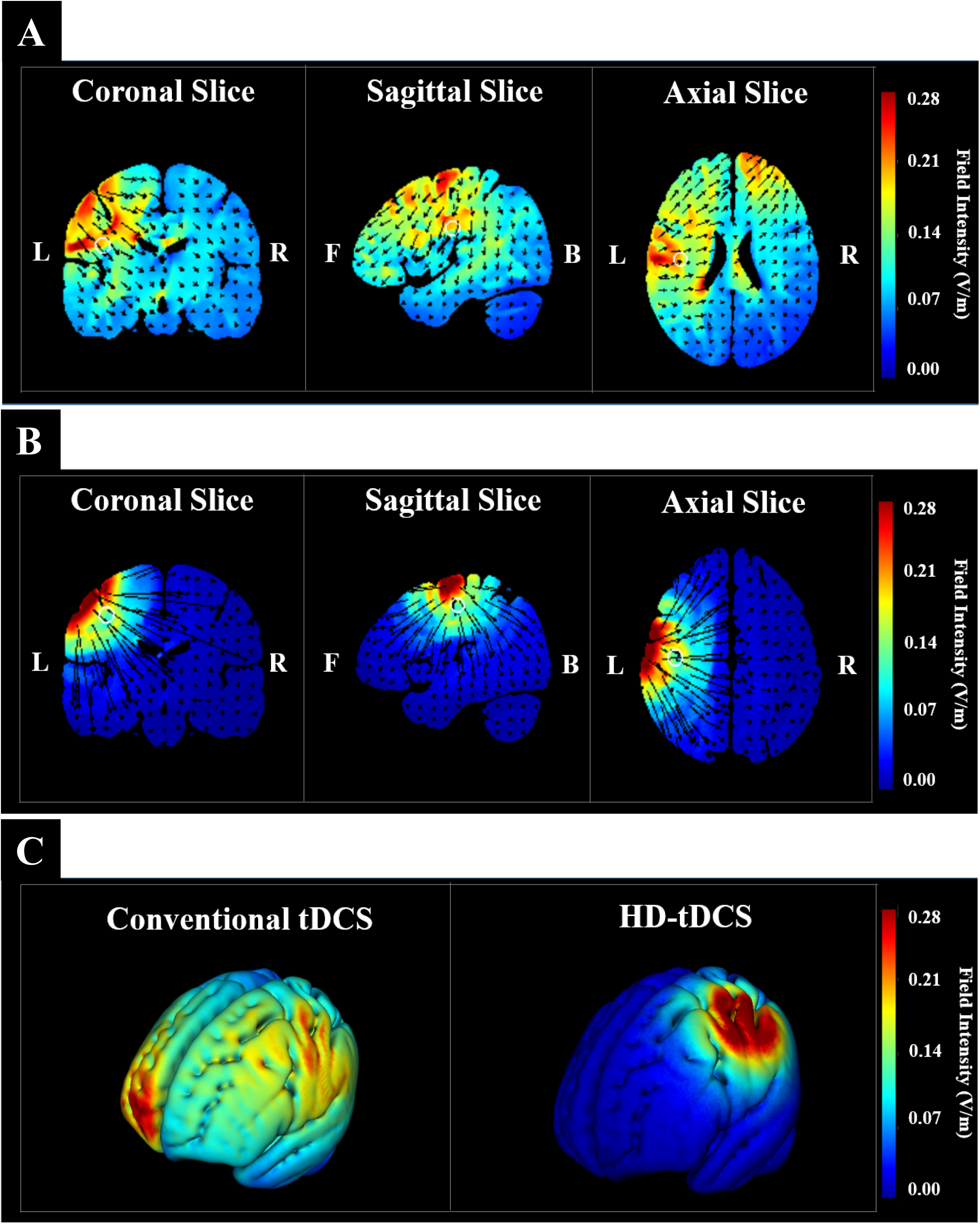
(a) Conventional a-tDCS electrode montage. Digital reconstruction displaying coronal, sagittal and axial views of the dispersion of electric current from the active electrode over M1 to the return electrode on the contralateral supraorbital region (b) HD a-tDCS electrode montage. Digital reconstruction displaying coronal, sagittal and axial views of the dispersion of electric current from the active electrode over M1 to the four return electrodes positions in a ring formation 3.5cm from M1. Four surrounding return electrode spaced 3.5cm from the central active electrode. (c) Digital reconstruction of the brain outling the differences in electric current dispersion between conventional tDCS and HD-tDCS. For all digital reconstructions, high EF intensity is displayed in red while lowest EF intensity is displayed in blue. Images adapted from Soterix Medical Inc. HD-explore software https://soterixmedical.com/research/software/hd-explore (Soterix Medical Inc, 2019)

One investigation was conducted by Kuo et al., (2013). Changes in CSE, as measured by the amplitude of transcranial magnetic stimulation (TMS)-evoked motor evoked potential (MEP), were directly compared following anodal and cathodal conventional tDCS and HD-tDCS. Immediately following conventional anodal tDCS (a-tDCS), MEP amplitudes were significantly greater compared to HD-tDCS while at 30-minutes post a-tDCS MEP amplitudes were significantly greater following HD-tDCS compared to conventional tDCS. A similar delayed effect was reported following cathodal tDCS (c-tDCS) with MEP amplitudes significantly lesser 30-minutes following HD-tDCS compared to conventional tDCS. It was concluded that while magnitudes differed, as with conventional tDCS, the responses to HD-tDCS were polarity-dependent and investigations into changes in cortico-cortical excitability via assessment of intracortical inhibitory and facilitatory mechanisms should be conducted. The effect of HD-tDCS on cortico-cortical excitability or whether its increased focality properties influences intra-individual variability is yet to be investigated.

This study therefore aimed to investigate the effect both conventional and HD-tDCS electrode montages have on CSE, cortico-cortical excitability and intra-individual variability. We hypothesised that with previous reports of increased focality of HD-tDCS (Datta et al., 2008, 2009), the magnitude of change in CSE and cortico-cortical excitability would be greater in HD-tDCS compared to conventional tDCS. Additionally, we hypothesised that intra-individual variability would reduce following HD-tDCS when compared to conventional tDCS. The results of this study will help inform future studies on whether conventional or HD-tDCS are capable of reducing intra-individual variability and therefore which technique is most appropriate when investigating changes in CSE following tDCS in large sample sizes.

## Materials and Methods

### Study Design

A repeated measures randomised cross-over design was utilised. All participants provided written informed consent and ethics approval was granted by Monash University Human Ethics Research Committee.

### Participants

Fifteen healthy male volunteers with mean (±Standard Deviation–SD) age of 28.53±8.28 years attended four sessions. Female participants were excluded to control for potential effect of fluctuating estrogen and progesterone hormones throughout the menstrual cycle (Inghilleri et al., 2004; Smith et al., 2002; Zoghi et al., 2015). Participant handedness (12 Right-handed, 3 Left-handed) was determined by the Edinburgh Handedness Inventory (Oldfield, 1971). No participants reported neurological or psychological symptoms via the TMS adult safety-screening questionnaire (Keel et al., 2001).

### Electromyography (EMG)

Participants were seated in an adjustable chair with their dominant hand resting on a pillow. Participants’ skin was abraded and cleaned to minimise skin impedence (Gilmore & Meyers, 1983). MEPs were recorded from the dominant first dorsal interosseous (FDI) muscle at rest via pre-gelled self-adhesive bipolar Ag/AgCl disposable surface electrodes with 2cm inter-electrode distance positioned over the muscle belly (Kendell et al., 2010) with a ground electrode placed over the ulna styloid process. EMG signals were filtered, amplified (10-500Hz x 1000) and sampled at 1000Hz. All data were recorded on PC software (LabChart™, ADInstruments, Australia) via a laboratory analogue-digital interface (Powerlab, ADInstruments, Australia).

### Measurement of CSE and cortico-cortical excitability by TMS

Single and paired-pulse stimuli were delivered by a 70mm figure-of-eight magnetic coil (Magstim Company Limited, UK). The coil was held over the dominant primary motor cortex (M1) and oriented at 45° to the midline and tangential to the scalp to ensure current flowed antero-posteriorly (Rossini & Rossi, 1998). The coil was manually positioned over M1 to determine the cortical area and coil orientation that elicited the greatest MEP in the FDI muscle. Considered the hotspot, this spot was marked on the scalp with a marker to maximise coil placement consistency through the testing session.

Resting motor threshold (RMT) was then determined. A percentage of the TMS device maximal stimulator output (MSO), RMT was defined as the minimum stimulus intensity required to evoke an MEP peak-to-peak amplitude greater than 50µV in the target muscle in 5/10 consecutive stimuli (Devanne et al., 2006). Set to 50% of MSO, the TMS device was adjusted by 1-2% intervals until the elicited MEP peak-to-peak amplitude matched as above (Rothwell et al., 1999). The RMT was used to calculate the conditioning pulse for paired-pulse stimuli to measure cortico-cortical excitability.

The test intensity was then determined. For CSE measurement, the test intensity was the percentage of the TMS device MSO required to evoke an MEP peak-to-peak amplitude of approximately 1mV. Set to 50% of MSO, the TMS device was again adjusted in 1-2% intervals until on average the MEP peak-to-peak amplitude was approximately 1mV. The resultant test intensity was used for single-pulse and paired-pulse stimuli to measure CSE and cortico-cortical excitability. Determining the hotspot, RMT and test intensity as above were repeated for each of the four testing sessions.

### Outcome Measures

Single-pulse TMS-evoked MEPs were used as an index of CSE. Twenty-five single-pulse MEPs, separated by a 6sec inter-stimulus interval (ISI) at the test intensity, were recorded with mean peak-to-peak amplitude taken as a measure of CSE. Paired-pulse TMS-evoked MEPs were used to assess cortico-cortical excitability. Short-interval intracortical inhibition (SICI) and Intracortical facilitation (ICF) were assessed by delivering a subthreshold conditioning pulse (80% RMT) followed by a suprathreshold pulse at the test intensity. The paired-pulses were separated by inter-pulse intervals (IPI) of 3ms for SICI and 10ms for ICF (Di Pino et al., 2014; Kujirai et al., 1993). Fifty paired-pulse MEPs were delivered, 25 with 3ms IPI and 25 with 10ms IPI, in a randomised order, with an ISI of 6sec. Mean MEP peak-to-peak amplitude of the 25 paired-pulses were calculated and expressed as a percentage of the single-pulse mean MEP peak-to-peak amplitude (Di Pino et al., 2014; Kujirai et al., 1993). These were considered an index of intracortical inhibition and ICF of M1 (Di Pino et al., 2014; Kujirai et al., 1993).

Intra-individual variability was measured by single-pulse MEP standardised z-value SDs as previously utilised (Fernandez et al., 2017; Pellegrini et al., 2018c). For each participant (n=15), intervention (conventional a-tDCS, conventional c-tDCS, HD a-tDCS, HD c-tDCS) and time-point (baseline, 0-minutes, 30-minutes), z-values were calculated from single-pulse MEP data. Each individual single-pulse MEP peak-to-peak amplitude was compared to the grand MEP mean and SD for that particular intervention and time-point using the following formula: z-value=[(individual MEP–grand MEP mean)/grand MEP SD]. For each participant, 25 z-values were generated for each intervention and time-point, of which the SD was then calculated. This was considered a representation of the degree a each individual MEP value for a particular participant varied from the grand MEP mean for a particular intervention and time-point (Pellegrini et al., 2018c) and was considered an index of intra-individual variability.

### Transcranial direct current stimulation

Conventional tDCS and HD-tDCS were delivered via the MXN-9 HD-tDCS Stimulator (Soterix Medical Inc, USA). Current intensity was 1mA and was applied for 10-minutes with 30-second fade-in and fade-out periods.

#### Conventional

Conventional tDCS was administered via two rectangular saline-soaked sponge electrodes fixed to the scalp via velcro straps. The active electrode (4×6cm) was placed over the dominant M1 while the return electrode (5×7cm) was placed over the contralateral supraorbit region. The larger size of the return electrode minimised side-effects and reduced current density under the return electrode, focusing it under the active electrode (M. A. Nitsche et al., 2007).

#### High-Definition

HD-tDCS was administered via a 4×1 ring electrode montage Kuo et al., (2013). A central active electrode was placed over the cortical area for FDI with four indentical return electrodes positioned in a ring formation 3.5cm from the centre of the active ring electrode. Each electrode was circular and 12 millimetres in diameter. The ring electrodes were stabilised by custom plastic holders with conducting transmission gel filling the space between electrode and scalp (Soterix Medical Inc, USA).

### Procedure

The four interventions (conventional a-tDCS, conventional c-tDCS, HD a-tDCS, HD c-tDCS) were conducted in randomised order. Data collection order was consistent across all four testing sessions and was collected by a single assessor. Participants refrained from drinking alcohol, coffee or energy drinks at least 12 hours prior to participating to avoid effects of ethanol (Conte et al., 2008; Roberto et al., 2006) and caffeine (Cerqueira et al., 2006; Concerto et al., 2017) on CSE and cortico-cortical excitability. A minimum 48-hour washing-out period eliminated risk of carry-over effects (Michael A. Nitsche et al., 2008). Each session was conducted at a similar time-of-day to minimise cortisol diurnal effects on CSE and cortico-cortical excitability (M. V. Sale et al., 2008; Martin V. Sale et al., 2007).

Once participants were seated and RMT and test intensity were determined, baseline outcome measures were recorded. One of the four tDCS interventions was then delivered. Immediately following this, outcome measures were recorded at 0-minutes post-tDCS. Single-pulse MEPs were recorded at the original baseline test intensity. RMT and test intensity to elicit ∼1mV were then re-determined and used to record the paired-pulse MEPs for SICI and ICF calculation. At 30-minutes post-tDCS, single-pulse and paired-pulse MEPs were recorded to investigate lasting effects (figure 2).

**Figure 2.**
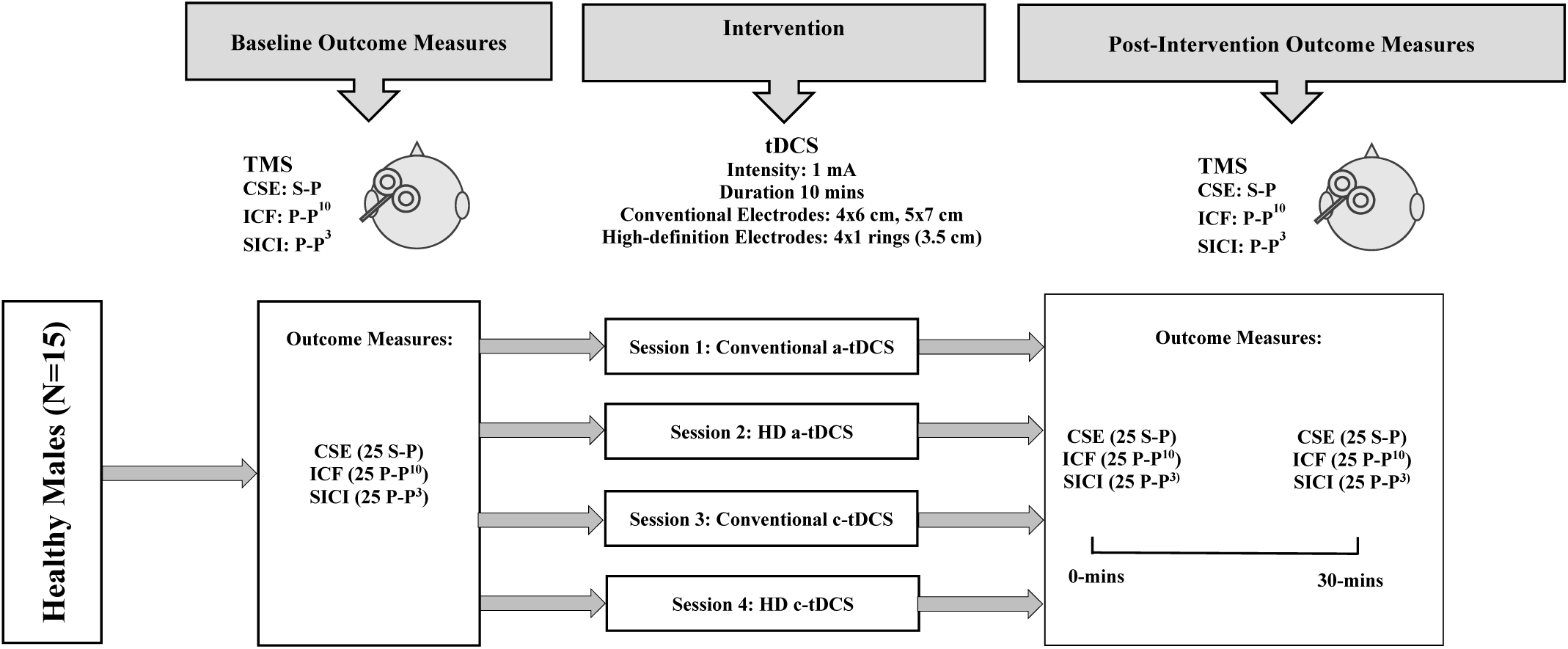
Experimental design. 25 Single and 50 paired-pulse TMS MEPs delivered to the M1 hotspot for FDI. Both anodal and cathodal conventional and HD tDCS delivered before re-assessing outcome measures.

### Data management and statistical analyses

Statistical analyses were conducted using SPSS (Version 22.0, IL, USA). All outcome measures (MEP, SICI, ICF and SDs of standardised z-values) were assessed for each intervention (conventional a-tDCS, conventional c-tDCS, HD a-tDCS, HD c-tDCS) and time-point (baseline, 0-minutes, 30-minutes). Tests for normality were conducted for each outcome measure using the Shapiro-Wilk test. If data violated normality, data were log-transformed to correct for data skewness as previously conducted in TMS studies (Hinder et al., 2014; Wassermann, 2002; Wiethoff et al., 2014). QQ and residual plots were also generated to assess data normality.

Two two-way repeated measures analysis of variance (RM-ANOVA) investigated the effect of time and each intervention on CSE and intra-individual variability. Post-hoc comparisons using the bonferroni correction were conducted where indicated. Conventional a-tDCS was compared to HD a-tDCS, while conventional c-tDCS was compared to HD c-tDCS. Intervention (two levels) and time (three levels) were the independent factors. Mauchly’s sphericity test investigated whether the variances of the differences between time-points and interventions were equal. Significance levels of p>0.05 indicated variances of the difference were equal and sphericity was assumed.

Significance levels of p<0.05 indicated variances were unequal, sphericity was not assumed and Greenhouse-Geisser corrected significance levels were used. When appropriate, one-way RM-ANOVA were performed to test whether post-tDCS CSE and intra-individual variability were significantly different from baseline and whether there were significant differences between interventions at each time-point. RM-ANOVA were performed at 5% significance levels (α=0.05).

To investigate the effect of intervention and time on cortico-cortical excitability (SICI and ICF), two two-way RM-ANOVA were conducted on log-transformed data to correct for skewness (Hinder et al., 2014; Wassermann, 2002; Wiethoff et al., 2014) with post-hoc comparisons via Bonferroni corrections. Independent factors were intervention (two levels) and time (three levels). Mauchly’s test for sphericity was again used and RM-ANOVA’s were performed at 5% significance level (α=0.05).

To avoid cumulative effect of changes in CSE and cortico-cortical excitability in repeated measures study designs (Alonzo et al., 2012; Gálvez et al., 2013; Michael A. Nitsche et al., 2008), between-session reliability was assessed using a two-way RM-ANOVA. For each outcome measure, baseline values from each session were used assessing the main effect of intervention on within-subjects reliability. Therefore reporting lack of carry-over effects and level of agreement between sessions (Portney & Watkins, 2000). Single assessor reliability of the TMS device operator has been previously reported (Pellegrini et al., 2018c).

## Results

All participants completed all four sessions with mean (±SD) intervals between each session 7.34±6.32 days. Mean RMT and test intensity were 36% (36.11±7.47) and 44% (43.95±8.86) of the TMS device MSO respectively.

### Tests for Normality

Shapiro-Wilk tests revealed single-pulse MEP peak-to-peak amplitude and standardised z-value SDs data were normally distributed for all tDCS interventions and time-points (p>0.05). Supporting this, a-tDCS and c-tDCS MEP peak-to-peak amplitude QQ plots (figure 3a, 4a) appear linear and residual plots (figure 3b, 4b) display random dispersion, suggesting MEP data do not violate normality. Tests for normality for SICI and ICF revealed data violated normality (p<0.05).

**Figure 3.**
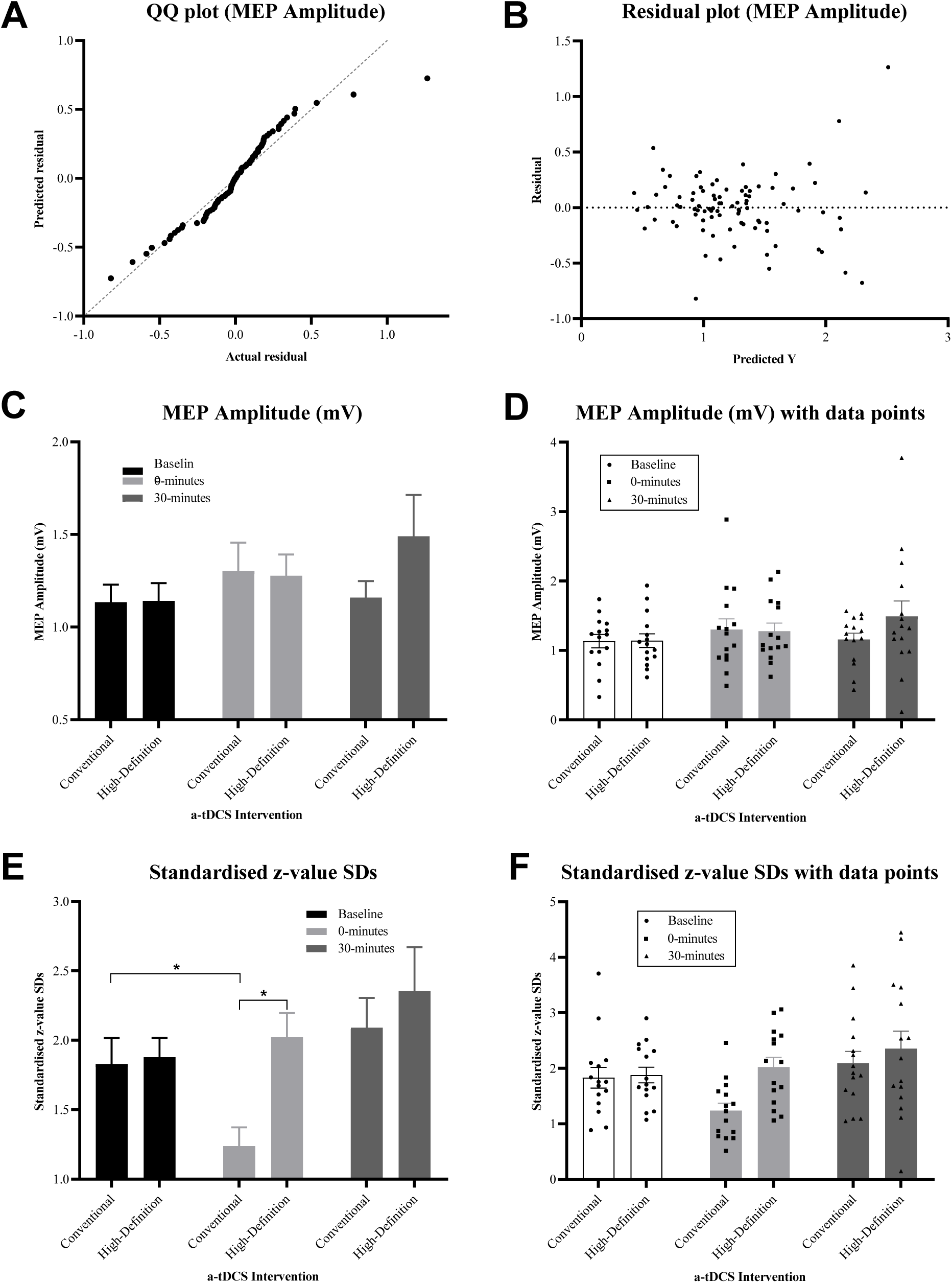
Anodal-tDCS (a) a-tDCS MEP amplitude Q-Q probability plot (b) a-tDCS MEP amplitude residual plot (c) MEP amplitude at each time-point following conventional a-tDCS and HD a-tDCS. (d) MEP amplitude at each time-point following conventional a-tDCS and HD a-tDCS with individual data points. (e) Intra-individual variability as represented by standardised z-value SDs at each time-point following conventional and HD a-tDCS.*denote significant differences between baseline and 0-minutes for conventional a-tDCS (p<0.01), and between conventional a-tDCS and HD a-tDCS at 0-minutes post-intervention (p<0.01) (f) Standardised z-value SDs at each time-point with individual data points. Error bars denote one SEM.

**Figure 4.**
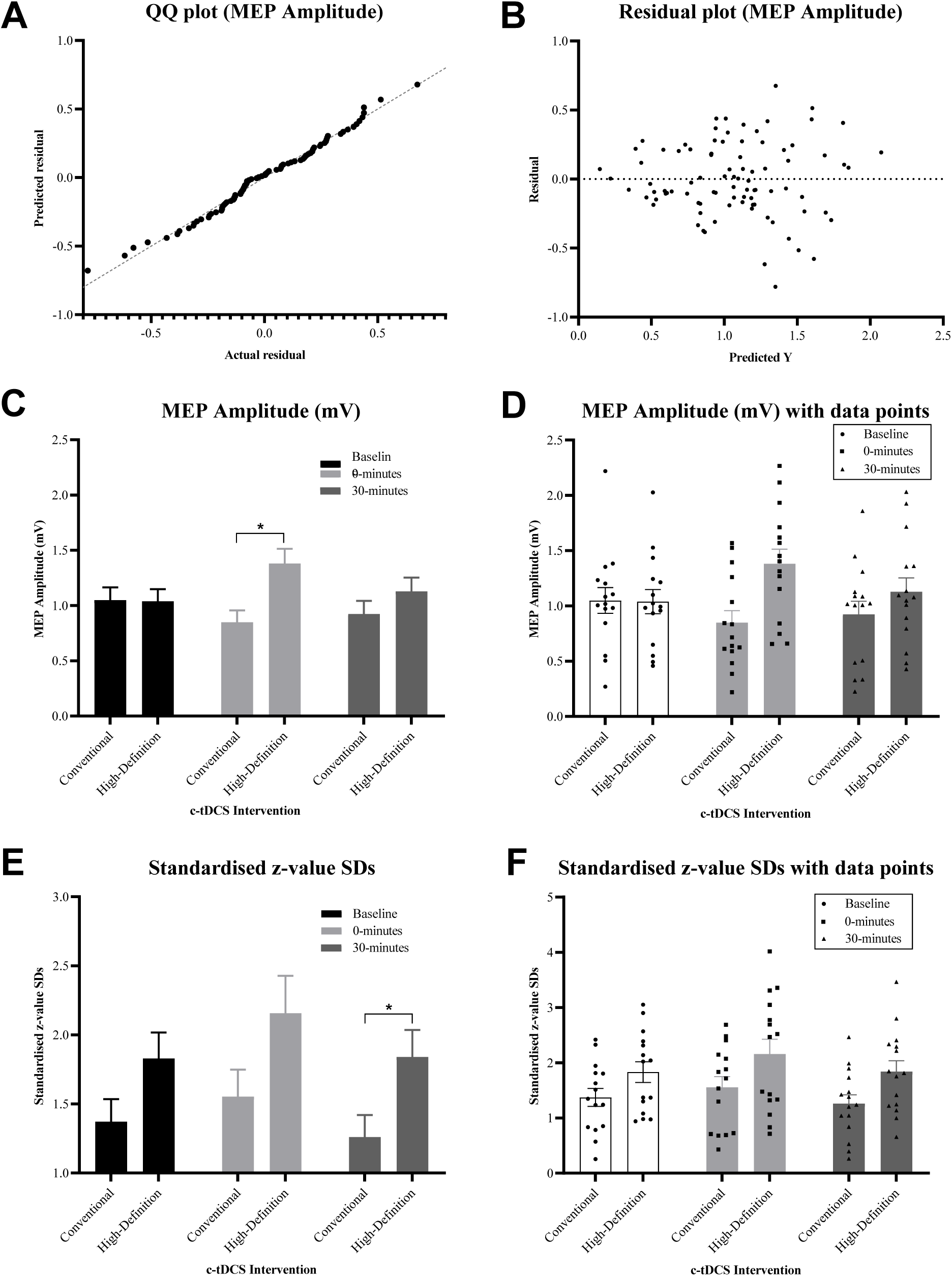
Cathodal-tDCS (a) c-tDCS MEP amplitude Q-Q probability plot (b) c-tDCS MEP amplitude residual plot (c) MEP amplitude at each time-point following conventional c-tDCS and HD c-tDCS. (d) MEP amplitude at each time-point following conventional c-tDCS and HD c-tDCS with individual data points. (e) Intra-individual variability as represented by standardised z-value SDs at each time-point following conventional and HD c-tDCS.*denote significant difference between conventional c-tDCS and HD c-tDCS at 30-minutes post-tDCS (p=0.010) (f) Standardised z-value SDs at each time-point with individual data points. Error bars denote one SEM.

### Carry-Over Effects and Level of Agreement

RM-ANOVA did not reveal significant differences in baseline values between interventions for MEP peak-to-peak amplitude (p=1.000), standardised z-value SDs (p=0.297), log-transformed SICI (p=1.000) and log-transformed ICF (p=1.000), indicating no carry-over effects between experimental sessions and single assessor intra-rater reliability.

### Comparison of conventional a-tDCS with HD a-tDCS

#### Corticospinal Excitability

Two-way RM-ANOVA showed no significant main effect of time (p=0.124), intervention (p=0.431) or time*intervention interaction (p=0.127). Mean baseline values for MEP peak-to-peak amplitude were 1.134±0.369mV and 1.141±0.376mV for conventional a-tDCS and HD a-tDCS respectively. MEP amplitude increased to 1.302±0.599mV 0-minutes following conventional a-tDCS (p>0.05) and 1.279±0.449mV following HD a-tDCS (p>0.05). MEP amplitude reduced toward baseline to 1.159±0.347mV at 30-minutes following conventional a-tDCS (p>0.05) while continued to increase to 1.490±0.866mV following HD a-tDCS (p>0.05) (Figures 3c, 3d and Table 1).

**Table 1.**
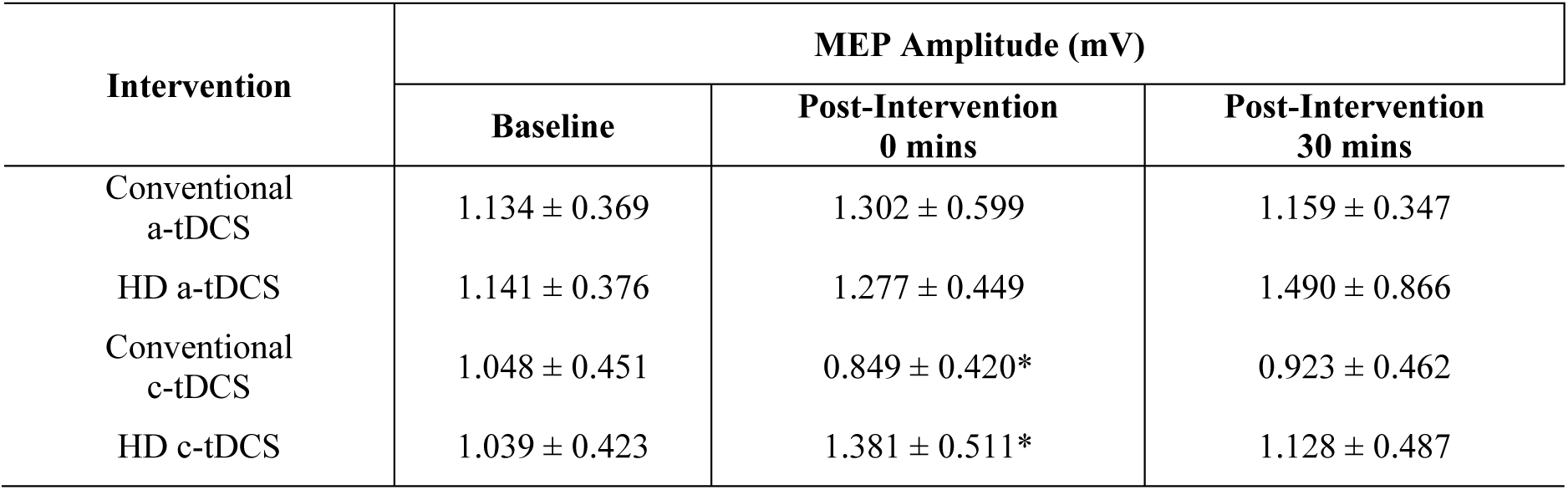
Corticospinal Excitability. Mean (±SD) MEP amplitude (mV) for each intervention and time-point.*denote significant difference in MEP amplitude between conventional c-tDCS and HD c-tDCS at 0-minutes post-tDCS (p=0.035)

| Intervention | MEP Amplitude (mV) |  |  |
| --- | --- | --- | --- |
|  | Baseline | Post-Intervention<br>0 mins | Post-Intervention<br>30 mins |
| Conventional<br>a-tDCS | 1.134 ± 0.369 | 1.302 ± 0.599 | 1.159 ± 0.347 |
| HD a-tDCS | 1.141 ± 0.376 | 1.277 ± 0.449 | 1.490 ± 0.866 |
| Conventional<br>c-tDCS | 1.048 ± 0.451 | 0.849 ± 0.420* | 0.923 ± 0.462 |
| HD c-tDCS | 1.039 ± 0.423 | 1.381 ± 0.511* | 1.128 ± 0.487 |

#### Response Variability

Two-way RM-ANOVA showed significant main effects of time (p<0.01), intervention (p=0.047) and time*intervention interaction (p=0.022). Mean baseline values for standardised z-value SDs as an index of intra-individual variability were 1.830±0.724 and 1.188±0.545 for conventional a-tDCS and HD a-tDCS respectively. Intra-individual variability reduced to 1.238±0.599 0-minutes following conventional a-tDCS (p<0.01) and increased to 2.021±0.675 following HD a-tDCS (p>0.05). Intra-individual variability increased to 2.091±0.830 30-minutes following conventional a-tDCS (p>0.05) and continued to increase to 2.354±1.230 following HD a-tDCS (p>0.05). Post-hoc analyses revealed no significant difference at baseline (p>0.05), while at 0-minutes post-tDCS intra-individual variability was significantly lesser following conventional a-tDCS compared to HD a-tDCS (p<0.01). There were no significant differences in intra-individual variability between conventional a-tDCS and HD a-tDCS at 30-minutes post-tDCS (p>0.05) (Figures 3e, 3f and Table 2).

**Table 2.** Intra-Individual Variability. Mean (±SD) Standardised z-value SDs for each intervention and time-point.*denote significant difference between conventional a-tDCS and HD a-tDCS at 0-minutes post-tDCS (p<0.01) and between conventional c-tDCS and HD c-tDCS at 30-minutes post-tDCS (p=0.010).^†^denote significant difference between baseline and 0-minutes for conventional a-tDCS (p<0.01).

| Intervention | Average standardised z-value SDs |  |  |
| --- | --- | --- | --- |
|  | Baseline | Post-Intervention<br>0 mins | Post-Intervention<br>30 mins |
| Conventional a-tDCS | $1.830 \pm 0.724^{\dagger}$ | $1.238 \pm 0.522^{\dagger*}$ | $2.091 \pm 0.830$ |
| HD a-tDCS | $1.877 \pm 0.545$ | $2.021 \pm 0.675^*$ | $2.353 \pm 1.230$ |
| Conventional c-tDCS | $1.371 \pm 0.634$ | $1.553 \pm 0.754$ | $1.261 \pm 0.612^*$ |
| HD c-tDCS | $1.830 \pm 0.729$ | $2.157 \pm 1.047$ | $1.840 \pm 0.757^*$ |

#### Cortico-cortical Excitability

Two-way RM-ANOVA on log-transformed SICI and ICF data revealed no significant main effects of time (p>0.05), intervention (p>0.05) or time*intervention interaction (p>0.05) (Tables 3-5).

**Table 3.** Cortico-cortical Excitability. Raw Mean (±SD) SICI data for each intervention and time-point.

| Intervention | SICI (% of single-pulse MEPs) |  |  |
| --- | --- | --- | --- |
|  | Baseline | Post-Intervention<br>0 mins | Post-Intervention<br>30 mins |
| Conventional<br>a-tDCS | 47.81 ± 27.09% | 44.67 ± 23.16% | 53.63 ± 32.49% |
| HD a-tDCS | 40.33 ± 23.67% | 42.42 ± 27.80% | 43.62 ± 34.98% |
| Conventional<br>c-tDCS | 37.76 ± 37.08% | 40.03 ± 35.80% | 56.32 ± 40.22% |
| HD c-tDCS | 47.26 ± 39.96% | 42.03 ± 27.23% | 51.09 ± 36.54% |

**Table 4.** Cortico-cortical Excitability. Raw Mean (±SD) ICF data for each intervention and time-point.

| Intervention | ICF (% of single-pulse MEPs) |  |  |
| --- | --- | --- | --- |
|  | Baseline | Post-Intervention<br>0 mins | Post-Intervention<br>30 mins |
| Conventional a-tDCS | 122.39 ± 52.28% | 123.63 ± 62.58% | 109.33 ± 50.43% |
| HD a-tDCS | 113.51 ± 53.75% | 124.88 ± 89.73% | 107.21 ± 44.12% |
| Conventional c-tDCS | 106.50 ± 55.34% | 103.78 ± 53.07% | 134.71 ± 57.97% |
| HD c-tDCS | 114.73 ± 59.70% | 108.05 ± 47.77% | 142.38 ± 63.51% |

**Table 5.** Cortico-cortical Excitability. Mean (±SD) of log-transformed SICI and ICF (%) for each intervention and time-point.

| Intervention | SICI (log-transformed) |  |  | ICF (log-transformed) |  |  |
| --- | --- | --- | --- | --- | --- | --- |
|  | Baseline | Post-Intervention<br>0 mins | Post-Intervention<br>30 mins | Baseline | Post-Intervention<br>0 mins | Post-Intervention<br>30 mins |
| Conventional a-tDCS | -0.395 ± 0.271 | -0.401 ± 0.221 | -0.348 ± 0.277 | .047 ± 0.198 | 0.02 ± 0.287 | -0.019 ± 0.252 |
| HD a-tDCS | -0.474 ± 0.283 | -0.451 ± 0.277 | -0.485 ± 0.337 | -0.014 ± 0.294 | 0.013 ± 0.285 | -0.002 ± 0.174 |
| Conventional c-tDCS | -0.598 ± 0.409 | -0.519 ± 0.324 | -0.370 ± 0.359 | -0.021 ± 0.209 | -0.032 ± 0.209 | 0.099 ± 0.162 |
| HD c-tDCS | -0.444 ± 0.328 | -0.496 ± 0.373 | -0.376 ± 0.275 | 0.008 ± 0.222 | -0.027 ± 0.276 | 0.098 ± 0.268 |

### Comparison of conventional c-tDCS with HD c-tDCS

#### Corticospinal Excitability

Two-way RM-ANOVA showed no significant main effect of time (p=0.639) but a significant main effect for intervention (p=0.037) and time*intervention interaction (p<0.01). Mean baseline values for MEP peak-to-peak amplitude were 1.048±0.451mV and 1.039±0.423mV for conventional c-tDCS and HD c-tDCS respectively. MEP amplitude reduced 0.849±0.420mV 0-minutes following conventional c-tDCS (p>0.05), but increased to 1.381±0.511mV following HD c-tDCS (p=0.047)). MEP amplitude increased toward baseline to 0.923±0.462mV at 30-minutes following conventional c-tDCS (p>0.05), while reduced toward baseline to 1.128±0.487mV following HD c-tDCS (p>0.05). Post-hoc analyses revealed no significant differences at baseline (p>0.05), while at 0-minutes post-tDCS MEP amplitude was significantly greater following HD c-tDCS (p<0.01). There were no significant differences at 30-minutes post-tDCS (p>0.05) (Figure 4c, 4d and Table 1).

#### Response Variability

Two-way RM-ANOVA showed no significant main effect of time (p>0.05), a significant main effect of intervention (p=0.015) but no significant time*intervention interaction (p=0.749). Mean baseline values for intra-individual variability were 1.371±0.634 and 1.830±0.792 for conventional c-tDCS and HD c-tDCS respectively. Intra-individual variability increased to 1.553±0.754 0-minutes following conventional c-tDCS (p>0.05) and increased to 2.157±1.047 following HD c-tDCS (p>0.05). Intra-individual variability then reduced to 1.261±0.612 30-minutes following conventional c-tDCS (p>0.05) and reduced to 1.840±0.757 following HD c-tDCS (p>0.05). Post-hoc analyses revealed no significant differences at baseline or 0-minutes post-tDCS (p>0.05). At 30-minutes post-tDCS intra-individual variability was lesser following conventional c-tDCS compared to HD c-tDCS (p<0.01) (figure 4e, 4f and table 2).

#### Cortico-cortical Excitability

Two-way RM-ANOVA on log-transformed SICI and ICF data revealed no significant main effect of time (p>0.05), intervention (p>0.05) or time*intervention interaction (p>0.05) (tables 4 and 5).

## Discussion

This study compared the effect of conventional tDCS and HD-tDCS on CSE, cortico-cortical excitability and intra-individual variability. It was hypothesised that due to increased focality properties of HD-tDCS, compared to conventional tDCS the magnitude of change in CSE and cortico-cortical excitability would be greater while intra-individual variability would be reduced. No significant differences in CSE were reported between both a-tDCS interventions, but significant differences were reported between c-tDCS interventions. No significant changes in cortico-cortical excitability were reported. Lastly, significant differences in intra-individual variability were reported between a-tDCS interventions as well as between c-tDCS interventions.

### Comparison of conventional a-tDCS with HD a-tDCS

It was hypothesised that due to its increased focality properties, the magnitude of increase in CSE and change in cortico-cortical excitability would be greater following HD a-tDCS compared to conventional a-tDCS. The findings of this study reported no significant differences in CSE and cortico-cortical excitability between a-tDCS interventions. This finding is supported by previously published studies reporting no significant overall increases in CSE, as measured by MEP peak-to-peak amplitude, following 10-minutes of conventional a-tDCS at 1mA current intensity (Chew et al., 2015; Tremblay et al., 2016). In contrast to our findings, while changes in CSE followed a similar pattern, Kuo et al., (2013) reported CSE was significantly greater immediately following conventional a-tDCS and significantly greater 30-minutes following HD a-tDCS. While the electrode montages and stimulation durations were comparable between the current study and Kuo et al., (2013), differences in current intensity may have contributed to the differing results, with Kuo et al., (2013) appliying a current intensity of 2mA.

A current intensity of 1mA was selected for this study to maintain its consistency with recently published conventional tDCS studies with large sample sizes (Chew et al., 2015; Tremblay et al., 2016) as well as to ensure current density was not too high causing discomfort to the participating individuals (Michael A. Nitsche et al., 2008). Discrepancies in our current findings may suggest that while the direction and pattern of change in CSE is similar to that of Kuo et al., (2013), our utilised tDCS current intensity of 1mA did not sufficiently influence the magnitude of EF focalised over the cortical area of interest to evoke statistically significant changes in CSE. No changes in CSE were supported by no observed significant differences in cortico-cortical excitability, as measured by SICI and ICF, between conventional a-tDCS and HD a-tDCS. Given the changes in CSE for both conventional a-tDCS and HD a-tDCS followed the same patterns previous studies, conduting head-to-head comparisons between these two difference electrode montages in larger sample sizes may provide additional insight into their different mechanisms of action under anodal conditions.

The results of this study do however highlight potential issues with the electrode montage for HD-tDCS. There is an inherent assumption that the programmed current intensity for the surrounding return electrodes will not influence the underlying cortical area. For HD a-tDCS, the active was electrode set at 1mA, while the surrounding return electrodes placed 3.5cm apart were programmed to -0.25mA, effectively generating c-tDCS conditions at 0.25mA under these cortical areas. With M1 receiving functional connections from multiple surrounding cortical areas including the prefrontal cortex and somatosensory cortex (Dum & Strick, 1991; He et al., 1993), it is reasonable to suggest that even at low current intensities, stimulation of these cortical areas may influence overall M1 excitability. In support of this, recently published work reported reductions in M1 excitability following c-tDCS of 0.3mA over the primary somatosensory cortex as well as the dorsolateral prefrontal cortex (DLPFC) (Vaseghi, Zoghi, & Jaberzadeh, 2015). While different stimulus durations were applied in the study (20-minutes) compared to the current study (10-minutes), it is still an example of the relationship and functional connectivities between M1 and surrounding cortical areas. Therefore, the finding of no immediate difference in CSE between conventional a-tDCS and HD a-tDCS may be explained by the -0.25mA inhibitory input from the surrounding cortical areas negating or overriding the focalised 1mA current intensity over M1. To maximise the utility of HD-tDCS, the findings of this study highlight issues with the electrode montage that require addressing.

### Comparison of conventional c-tDCS with HD c-tDCS

As with a-tDCS, it was hypothesised that again due to its increased focality properties, the magnitude of reduction in CSE and change in cortico-cortical excitability would be greater following HD c-tDCS compared to conventional c-tDCS. This study reported the novel finding of an increase in CSE following HD c-tDCS compared to a reduction in CSE following conventional c-tDCS. This excitatory response immediately following HD c-tDCS was significantly greater than the inhibitory response following conventional c-tDCS at the samet time-point (Figure 4c, 4d and Table 1).

This novel finding is not consistent with previous literature and again may be explained by the differences in electrode montages utilised. For HD c-tDCS, the active electrodes are the four surrounding electrodes each set with a current intensity of 0.25mA while the central electrode is the return electrode set to -1mA. It is possible that a current intensity of 0.25mA at the surrounding electrodes is sufficient to induce an excitatory response in the underlying cortical region that is in close enough proximity to the cortical area of interest at M1. To these authors’ knowledge, increases in CSE in response to tDCS at low current intensities such as 0.25mA has not been reported. Investigations into the effect a similar current intensity of 0.3mA tDCS have however previously been investigated (Bastani & Jaberzadeh, 2013). Following 10-minutes of conventional a-tDCS over M1 with a current intensity of 0.3mA, significant increases in CSE were reported (Bastani & Jaberzadeh, 2013). More specifically, Vaseghi, Zoghi, & Jaberzadeh, (2015) reported increases in CSE of M1 following 20-minutes of a-tDCS at 0.3mA fover the DLPFC. It was again suggested that as the prefrontal cortex delivers inputs to the premotor areas (Dum & Strick, 1991; He et al., 1993), a-tDCS of the DLPFC, even at small current intensities of 0.3mA, may activate the functional connections between DLPFC, premotor areas and M1, thus increasing the excitability of M1 (Vaseghi et al., 2015). Therefore, for HD c-tDCS, it may be plausible to suggest a current intensity of 0.25mA from the return electrode positioned approximately over the DLPCF and other cortical areas may override the inhibitory effect of -1mA over M1. These previous studies along with the results of this current study challenge previous reports that tDCS current intensities of at least 0.6mA are required to induce significant increases in CSE (Bastani & Jaberzadeh, 2013; M. A. Nitsche & Paulus, 2000b).

To support our initial hypotheses that conventional c-tDCS and HD c-tDCS would reduce CSE, it was also hypothesised that there would be reductions in cortico-cortical excitability as measured by increases in SICI. The novel finding of significantly greater CSE immediately following HD c-tDCS compared to conventional c-tDCS was not supported by any changes in SICI or ICF for either of the c-tDCS interventions. Through previous TMS and pharmacological studies, SICI is considered a measurement of the activity of particular M1 intracortical GABA-ergic interneurons acting on GABA-A receptors (Ziemann et al., 1996; Zoghi et al., 2015). With the plausible suggestions that for HD c-tDCS, a current intensity of 0.25mA at the return electrodes may be sufficient to limit the inhibitory mechanism of the active electrode over M1, small observed reductions in SICI immediately following HD c-tDCS may contribute to the overall increase in CSE. The fact these changes in SICI were not significant however do not shed light into mechanisms of action of HD c-tDCS on cortico-cortical excitability. Future investigations into the effect of HD-tDCS on cortico-cortical excitability via SICI and ICF should aim to do so in larger sample sizes to increase the likelihood of reporting significant changes in cortico-cortical excitability.

The novel results of this current study highlight current technical issues with HD-tDCS particularly under cathodal conditions. With the possibility that a current intensity of 0.25mA in surrounding return electrodes over cortical areas with functional connections to M1 is sufficient to induce an excitatory response, investigation into methodologies for reducing the return electrode current intensity are necessary. One such suggestion may be to increase the number of electrodes arranged in the ring formation. For example, if the number of electrodes arranged in the ring formation is increased to greater than the standard four, the current intensity will be reduced at each electrode to less than 0.25mA, thus reducing the risk of inducing an unintended excitatory response. Addressing these technical issues will facilitate investigation into the true effects of HD c-tDCS on changes in M1 CSE. To these authors’ knowledge, this is yet to be investigated.

### Response Variability

We hypothesised that intra-individual variability, as measured by SDs of standardised z-scores, would be reduced following HD-tDCS compared to conventional tDCS. To our knowledge, this is the first study to compare intra-individual variability between conventional tDCS and HD-tDCS and our novel findings suggest that response variability increases following HD-tDCS compared to conventional tDCS, not supporting our original hypotheses.

The current quantifying technique for intra-individual variability have only been used on a number of occasions (Fernandez et al., 2017; Pellegrini et al., 2018c). Calculating the SDs from standardised data (z-scores) can be considered a robust method of quantifying intra-individual variability for a given sample of individuals. By quantifying the degree each individual data point deviates from the grand average for a given sample of individuals (Pellegrini et al., 2018c), this technique is normalised to the group level compared to other previous techniques where calculations remain at the level of the individual.. The coefficient of variation (CV) is one such example. The CV (SD/meanx100) represents the range, or percentage variance that one SD of the data lies about the mean (Atkinson & Nevill, 1998). Its simplicity has led to use in NIBS and TMS literature (eg: (Biabani et al., 2018; Klein-Flugge et al., 2013; Sadnicka et al., 2013; Martin V. Sale et al., 2017; Temesi et al., 2017; Vassiliadis et al., 2018)).

It is likely that a number of techniques may provide useful insight into response variability, however direct comparisons between these two quantifying techniques are yet to be carried out in the tDCS literature. Given the importance of response variability, future large-scale studies investigating the effects of tDCS on intra-individual variability, and the factors that may contribute, should quantify response variability with both techniques and conduct head-to-head comparisons to greater understand each technique’s ability to capture changes in intra-individual variability.

The significance of this current study lies in the novel finding that CSE following conventional tDCS appears less variable compared to HD-tDCS. This is despite reports of increased focality of HD-tDCS compared to conventional tDCS (Datta et al., 2008, 2009; Edwards et al., 2013). These results have implications for future studies aiming to investigate how tDCS modulates CSE and cortico-cortical excitability. Increases in intra-individual variability of CSE following HD-tDCS calls into question the validity of the observed changes in CSE following HD-tDCS, and whether they are true reflections of changes in CSE and cortico-cortical excitability. Therefore in its current form, it appears when addressing the ongoing issue of response variability in the tDCS literature, conventional tDCS may be better equipped compared to HD-tDCS.

### Limitations

The results of this current study were reported only on healthy males. While females were excluded to eliminate a potential source of variability due to fluctuating estrogen and progesterone levels (Inghilleri et al., 2004; Smith et al., 2002; Zoghi et al., 2015). It is unclear whether these results can be extrapolated to healthy young females. It is also unclear whether these current results can be extrapolated to older populations. Additionally, as outcome measures were only recorded at 30-minutes post-tDCS, the lasting effect of conventional tDCS and HD-tDCS on CSE, cortico-cortical excitability and response variability can not be inferred from the results of this current study.

### Future Directions

From a methodological perspective, future research should follow the growing trend of previous tDCS literature by investigating the effects of HD-tDCS on CSE, cortico-cortical excitability and response variability in large sample sizes similar to previous conventional tDCS electrode montage studies (Chew et al., 2015; Labruna et al., 2016; López-Alonso et al., 2014, 2015; Puri et al., 2015, 2016; Strube et al., 2015, 2016; Tremblay et al., 2016; Wiethoff et al., 2014). This will facilitate high powered investigation into the mechanisms of the more focalised HD-tDCS.

On a technical note, future research should refine the HD-tDCS electrode montage to investigate the effects a more concentrated electrode arrangement has on CSE and cortico-cortical excitability. As the current intensity of the surrounding return electrodes may play a role in the overall changes reported in CSE and cortico-cortical excitability, it should not be discounted when designing future HD-tDCS studies. Increasing the number of surround electrodes, may minimise the chance that activity in the cortical areas underlying the surround electrodes influences the overall effect of HD-tDCS, thus delivering more focalised current intensity to the targeted cortical area of interest as initially hypothesised with HD-tDCS electrode montages.

The significance of our findings and the implications for future research lie in the differences in intra-individual variability following both types of tDCS interventions. Currently reporting less intra-individual variability following conventional tDCS compared to HD-tDCS suggests that conventional tDCS may provide a more accurate representation of the true effects of tDCS on CSE and cortico-cortical excitability. This is particularly important for future large-scale studies aiming to investigate inter-individual variability and the question of why some individuals respond to tDCS as historically expected and why others do not. Future large-scale large sample size tDCS studies comparing conventional tDCS electrode montages with HD-tDCS refined as above, should aim to investigate whether tDCS electrode montage influences the number of individuals that respond as historically expected and the number that do not. At present however, given the differences in intra-individual variability between conventional tDCS and the current HD-tDCS, understanding the inherent anatomical and biological differences between individuals that influence inter-individual variability following tDCS may be better achieved utilising conventional tDCS electrode montages.

## Conclusions

This current study reported no significant changes in CSE following both conventional and HD a-tDCS, yet reported significantly greater CSE following HD c-tDCS compared to conventional c-tDCS. These findings were not supported by any significant changes in cortico-cortical excitability, as measured by SICI and ICF. This current study also reported the novel finding that conventional tDCS and HD-tDCS impact on intra-individual variability differently. Future studies aiming to explore the overall mechanism of action of tDCS on CSE and cortico-cortical excitability should build upon this current study and conduct head-to-head comparisons of these two electrode montages in large sample-sizes. At present however, with lower magnitudes of intra-individual variability after conventional tDCS compared to HD tDCS, it appears future large-scale investigations into the issue of response variability, conventional electrode montages provide more reliable and less variable results. With tDCS technology a popular means of investigating cortical plasticity, placing a focus on different techniques such as HD-tDCS, as in this study, are important steps to take to ultimately refine and optimise the delivery of tDCS. This is to ensure the technology can be consistently used to address important issues such as response variability and can progress to a meaningful application in neuropathological populations with the most predictable and reliable tDCS protocols.

## Abbreviations

a-tDCS: Anodal Transcranial Direct Current Stimulation
c-tDCS: Cathodal Transcranial Direct Current Stimulation
CSE: Corticospinal Excitability
CV: Coefficient of Variation
EMG: Electromyography
EF: Electric Field
FDI: First Dorsal Interossei
HD: High Definition
ICF: Intracortical Facilitation
IPI: Inter Pulse Interval
ISI: Inter Stimulus Interval
M1: Primary Motor Cortex
mA: Milliamperes
MEP: Motor Evoked Potential
ms: Millisecond
MSO: Maximal Stimulator Output
mV: Millivolt
NIBS: Non-Invasive Brain Stimulation
RM-ANOVA: Repeated Measures Analysis of Variance
RMT: Resting Motor Threshold
SD: Standard Deviation
SEM: Standard Error of the Mean
SICI: Short Interval Intracortical Inhibition
tDCS: Transcranial Direct Current Stimulation
TMS: Transcranial Magnetic Stimulation
µV: Microvolts

## Conflict of Interest Statement

This research did not receive any funding from agencies in the public, commercial, or not-for-profit sectors.

## Author Contributions

Conceived and designed study: MP, SJ; Carried out data collection: MP; Conducted the analysis: MP, SJ; Interpreted the findings: MP, SJ, MZ; Wrote the Manuscript: MP; Writing and editing of drafts: MP, SJ, MZ.

